# Effects of combined application of organic fertilizer and inorganic fertilizer on soil and yield of Gaodan grass

**DOI:** 10.1101/2024.12.19.629543

**Authors:** Zhixin Yi, Qiuxu Liu, Yuhui Chen, Yunqun Zhu, Yang Ji, Linxiang Lin

## Abstract

Using Sorghum bicolor × sudanense as test material, the effects of inorganic fertilizer and different proportions of organic fertilizer application on soil properties, yield and quality of sorghum were studied under conventional nitrogen application levels, aiming to provide data support for efficient production of green cycle forage. Five treatments were set up in the experiment, namely blank treatment (CK), conventional fertilization (HF), equivalent cow manure organic fertilizer (NF2), equivalent sheep manure organic fertilizer (YF2), 150% equivalent cow manure organic fertilizer (NF5) and 150% equivalent sheep manure organic fertilizer (YF5). In general, NF5 treatment was excellent in improving soil organic matter, enzyme activity and nutrient content, and the yield of Gaodan grass was significantly increased.

## 1. Introduction

At present, chemical fertilizer has been indispensable in the agricultural production process, which can quickly supplement soil nutrients. However, the utilization rate of applied chemical fertilizer is far below 50%^[1]^. Excessive and ineffective chemical fertilizer leads to excessive growth of plants, which are high and fine, prone to lodging and prone to diseases, and form the absorption antagonism of other nutrients. And even lower crop yields, which ultimately increase soil acidification and compaction, soil microecological imbalance ^[2-5]^. The decrease of soil fertility and crop yield reverse stimulate the increase of fertilizer use, and eventually form a vicious circle, resulting in soil and water pollution and ecological environment destruction ^[6]^. Fertilizer reduction substitution has become one of the hot spots in agriculture. Some studies have found that animal manure contains organic matter and nutrients such as nitrogen, phosphorus and potassium required by crops. Using animal manure instead of chemical fertilizer to return to the field can not only reduce animal manure pollution, but also make full use of the nutrients in animal manure to maintain crops. It has a positive effect on improving soil texture and maintaining soil microecology ^[7-10]^. Therefore, it is widely believed that organic fertilizer is far more effective than chemical fertilizer in improving soil fertility and improving soil texture. At the same time, organic fertilizer has a lasting effect on crop production and improving crop quality. It has become a trend to use organic fertilizer as raw material instead of chemical fertilizer ^[11-14]^.

## 2. Materials and methods

### 2.1 Test site condition

The experiment was conducted in Shanping Village, Lingyun Township, Bazhou District, Bazhong City, Sichuan Province, which belongs to the subtropical humid monsoon climate, with the annual average temperature of 16.2 °C and the annual total rainfall of 1719.3 mm. The soil in the experiment site was paddy soil, and the soil properties before the experiment were as follows: pH value 6.21, organic matter content 2.66%, total nitrogen content 1.25 g/kg, ammonia nitrogen content 14.04 mg/kg, nitrate nitrogen content 35.06 mg/kg, total phosphorus content 0.64 g/kg, total potassium content 12.78 g/kg, available phosphorus 116.16 mg/kg. Available potassium 62.39 mg/kg, urease activity 186.17 ug/d/g, phosphatase activity 15.72 umol/d/g, cation exchange capacity 21.45 cmol/kg.

### 2.2 Experiment design

The experiment was conducted in 2023 using a randomized block design with five treatments, each repeated three times: blank treatment (CK), conventional fertilization (HF), equivalent cow manure organic fertilizer (N2), equivalent sheep manure organic fertilizer (Y2), 150% equivalent cow manure organic fertilizer (N5), and 150% equivalent sheep manure organic fertilizer (Y5). The application amount of nitrogen, phosphorus and potassium in each treatment was the conventional application amount of Gaodan grass, namely N: 450 kg/hm^2^, P: 75 kg/hm^2^, K: 75 kg/hm^2^. Cattle manure organic fertilizer was fermented from Shau Ji Wan Ecological Farm in Chaba Town, Enyang District, Bazhong City (total nitrogen content 12.45 g/kg, total potassium content 10.34 g/kg, total phosphorus content 4.81 g/kg). Sheep manure organic fertilizer was fermented from sheep manure in Yuandingzi sheep Farm of Sichuan Beimu Nanjiang Yellow Sheep Group Co., LTD. (total nitrogen content 19.54 g/kg, total potassium content 8.74 g/kg, total phosphorus content 5.85 g/kg). Inorganic nitrogen, phosphorus and potassium are used to adjust, and the replacement ratio is calculated by the total amount of N. Among them, organic fertilizer, inorganic phosphate fertilizer and inorganic potassium fertilizer were all applied as base fertilizer at one time, and inorganic nitrogen fertilizer was applied three times, including base fertilizer, topdressing at jointing stage and topdressing after cutting, and the base-topdressing ratio was 1:1:1. Field management according to field production management. Each cell is 30 m^2^ (6 m×5 m), the cell interval is 1 m, and the surrounding protection line is 2 m. The national approved variety Shucao No. 4 Gaodan grass was independently selected by the Institute of Agricultural Resources and Environment of Sichuan Academy of Agricultural Sciences. The line spacing was 50 cm, and the sowing amount was 37.5kg /hm^2^.

### 2.3 Sample collection

Soil samples 0-20 cm soil samples were collected by random multi-point mixed sampling method. After air drying in a cool and dry place, the samples were screened by 2 mm. The plant samples were cut for the first time on June 23, 2023, and for the second time on August 20, 2023. Each cell removed the side lines and took the middle 8 lines for yield measurement. Take 5 kg of fresh grass from each plot to the laboratory for quality determination.

### 2.4 Index determination

Soil physical and chemical indexes: pH value, organic matter content, total nitrogen content, ammonia nitrogen content, nitrate nitrogen content, total phosphorus content, total potassium content, available phosphorus content, available potassium content, urease activity, phosphatase activity, cation exchange capacity.

Forage quality index: total nitrogen content, total phosphorus content, total potassium content, total calcium content, lignin content, starch content, soluble sugar content, neutral detergent fiber content, acid detergent fiber content, dry matter content, crude protein content, crude fat content, crude fiber content, ash content, nitrogen free extract content.

### 2.5 Statistical analysis

Mathematical Statistics and mapping were performed using Microsoft 2016, IBM Statistics SPSS 18.0 and Origin 2023 software for experimental data, and one-way analysis of variance was used for data analysis. The significance of multiple comparisons was tested by the method of least significant difference (LSD) (*p* = 0.05).

## 3. Results and discussion

### 3.1 Effect of different proportion of application on soil physicochemical properties

As shown in Figure 1, compared with the control group, soil pH value under HF and NF2 treatments significantly increased by 2.98% and 2.44%, respectively, while other treatments showed different degrees of decrease, among which YF2 treatment showed the most obvious decrease, reaching 2.00% (Fig. 1a). HF, NF2, NF5, YF2 and YF5 treatments can all effectively increase soil organic matter content, with the increase range ranging from 10.35% to 102.35%, among which the increase rate of NF5 treatment is the most significant, reaching 102.35% (Fig. 1b). Only NF2 treatment resulted in a 6.81% decrease in soil total nitrogen content, while all other treatments increased total nitrogen content, especially NF5 treatment, which increased by 69.47% (Fig. 1c). Similarly, NF2 treatment alone reduced soil ammonia nitrogen content by 13.80%, while other treatments all led to an increase in ammonia nitrogen content, especially NF5 and YF5 treatments, which increased by 76.92% and 93.76%, respectively (Fig. 1d). For soil nitrate nitrogen content, HF, NF2 and YF2 treatments all decreased, and the decrease was between 6.62% and 31.28%. In contrast, NF5 and YF5 treatments increased it by 1.67% and 6.03%, respectively (Fig. 1e). Only NF2 treatment resulted in a 3.74% decrease in soil total phosphorus content, while all other treatments resulted in an increase in total phosphorus content. NF5 and YF5 treatments had the largest increase, reaching 79.58% and 93.16%, respectively (Fig. 1f). Soil total potassium content under HF and YF2 treatments decreased by 16.56% and 16.87%, respectively, while other treatments increased total potassium content, and YF5 treatment showed the most obvious increase, reaching 27.74% (Fig. 1g). Only NF2 treatment resulted in a decrease in soil available phosphorus content.

**Figure 1.**
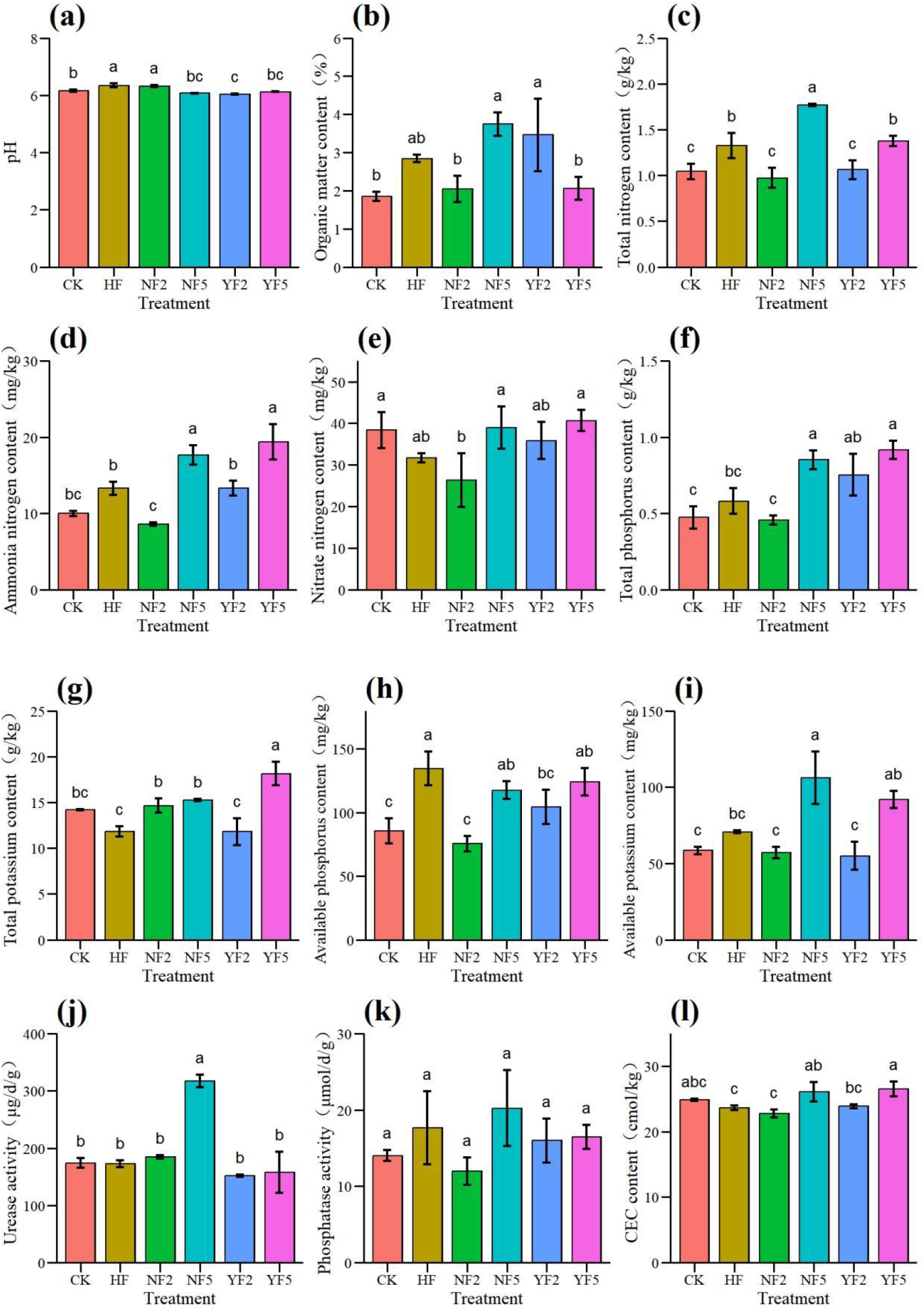
Effect of different proportion of application on soil physicochemical properties

### 3.2 Effect of different proportion of application on yield of Gaodan grass

According to Figure 2, compared with CK, the yield of Gaodan grass under HF, NF2, NF5, YF2 and YF5 treatments showed an increasing trend, with an increase range of 41.91% to 79.67%. Among them, the effect of HF, NF5 and YF2 treatment is particularly significant, and the yield increase is as high as 79.67%, 63.06% and 71.85%, respectively.

**Figure 2.**
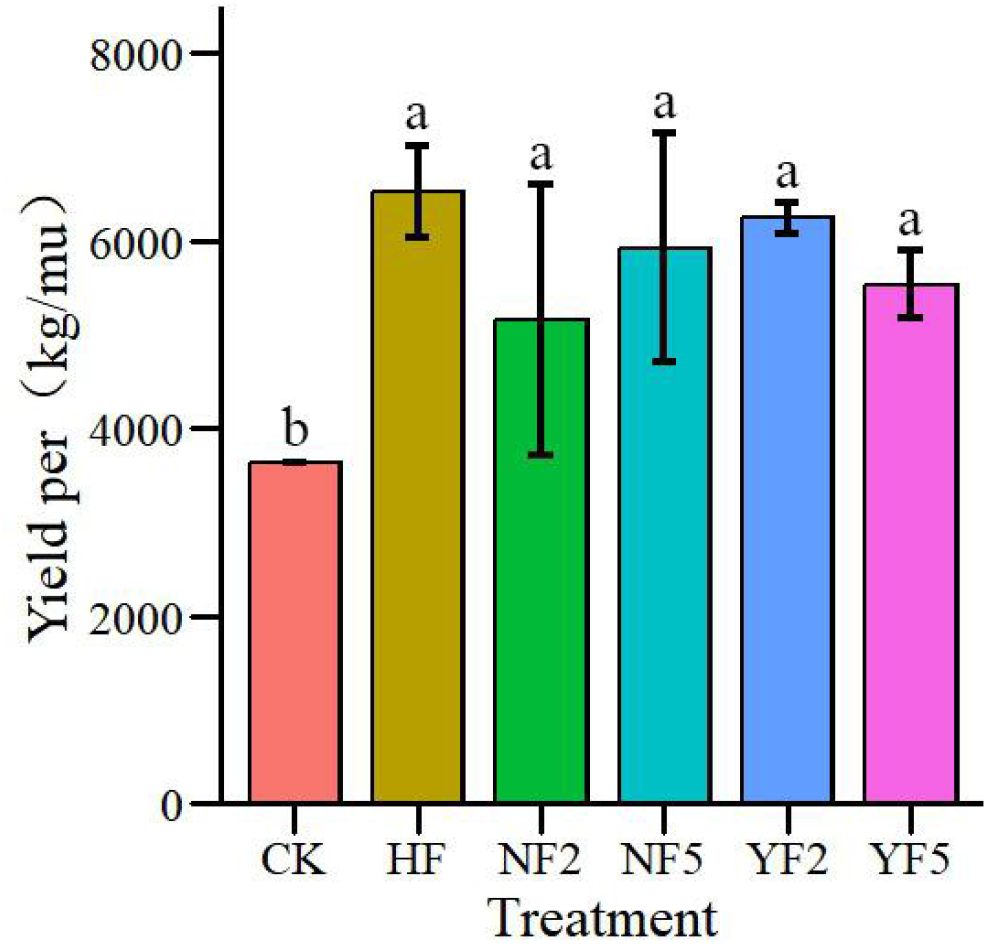
Effect of different proportion of application on yield of Gaodan grass

### 3.3 Effect of different proportion of application on quality of Gaodan grass

As shown in Figure 3, compared with CK, HF, NF2, NF5, YF2 and YF5 treatments all resulted in a decreasing trend of nitrogen content in Gaodan grass, ranging from 4.18% to 19.73%. Among them, the treatment effect of HF, NF2 and NF5 was particularly significant, with a decrease of 19.51%, 14.80% and 19.73%, respectively (Fig. 3a). In terms of total calcium content, compared with CK, all the above treatments significantly increased the total calcium content of Gaodan grass. In particular, HF and NF5 treatments showed a more significant improvement with increases of 42.25% and 53.56%, respectively (Fig. 3b). For the starch content, NF5 and YF2 treatments resulted in a decrease in the starch content of the grass, which was 13.67% and 4.74%, respectively. However, other treatments increased starch content, especially NF2 treatment, by 17.67% (Fig. 3c). In terms of soluble sugar content, NF5, YF2 and YF5 treatments resulted in a decrease in soluble sugar content of the grass, ranging from 0.97% to 8.68%. However, other treatments increased soluble sugar content, especially HF treatment, which increased by 23.18% (Fig. 3d). As for the neutral detergent fiber content, all treatments significantly increased the neutral detergent fiber content of Gaodan grass. Among them, HF treatment has the most obvious effect, with an increase of 11.18% (Fig. 3e). In terms of acid detergent fiber content, all treatments resulted in an increase in acid detergent fiber content of Gaodan grass, ranging from 2.64% to 9.15%, and the differences among treatments were not significant (Fig. 3f). As for the dry matter content, all the treatments increased the dry matter content of the grass, and the increase was between 15.55% and 20.97%.

**Figure 3.**
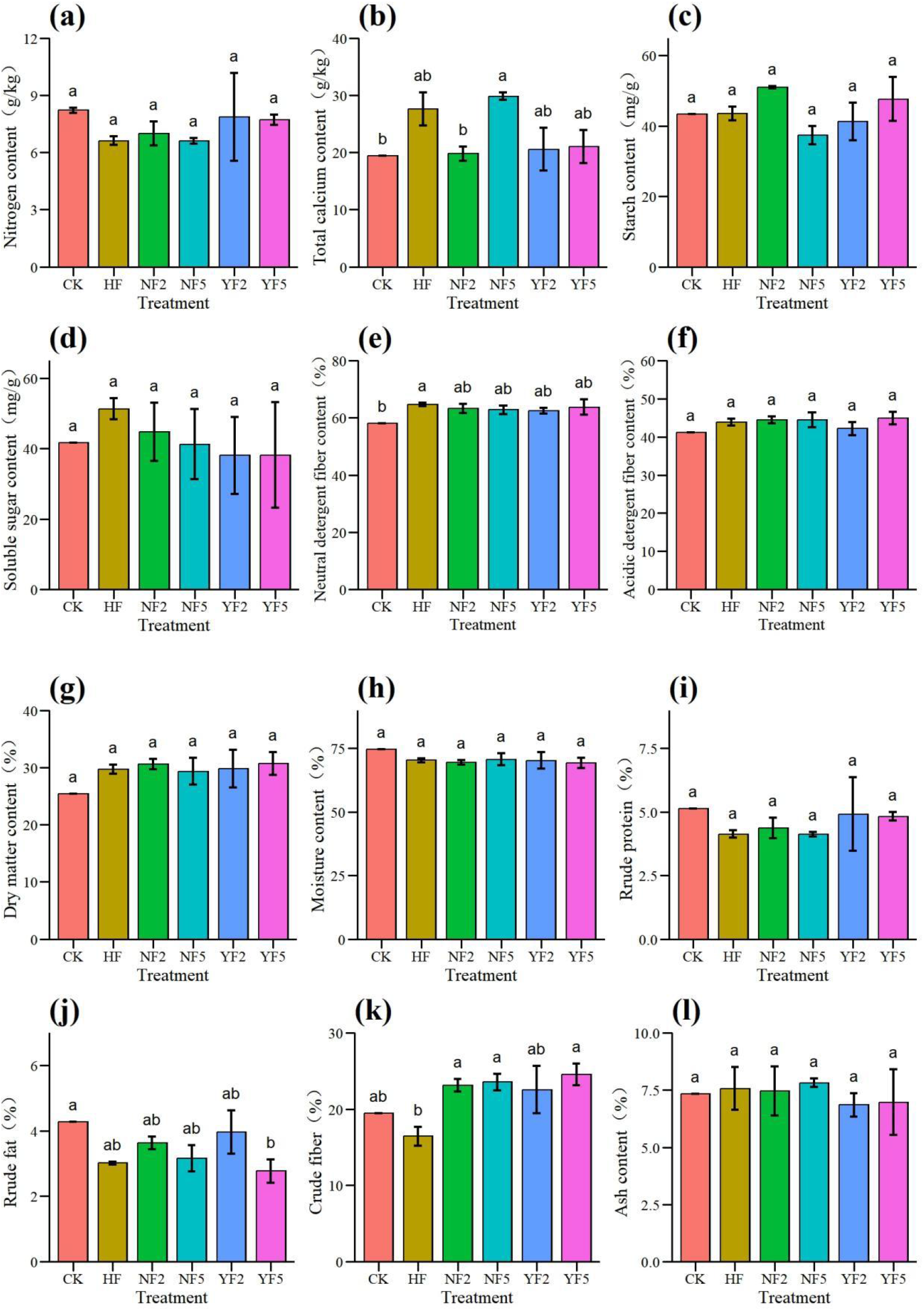
Effect of different proportion of application on quality of Gaodan grass

## 4. Conclusion

Different soil treatments had significant effects on soil indexes, yield and nutrient composition of the grass. HF and NF2 treatments increased soil pH value, while NF5 treatment showed excellent performance in improving soil organic matter, enzyme activity and nutrient content. However, NF2 treatment resulted in a decrease in nutrient content in some soils. The yield of Gaodan grass increased under all treatments, especially HF, NF5 and YF2 treatments. At the same time, the content of nitrogen, crude protein and crude fat decreased, while the content of total calcium, neutral detergent fiber, dry matter and nitrogen free extract increased. Among them, HF and NF5 treatment significantly increased the total calcium content, NF2 treatment increased the starch content, HF treatment significantly increased the soluble sugar and nitrogen free extract content. All treatments resulted in the decrease of water content, and the other components changed with different treatments, but the difference was not significant.

## Author Contributions

Zhixin Yi performed the bioinformatics analysis, performed the experiments and wrote the original manuscript. Qiuxu Liu performed the experiments and validation. Yuhui Chen performed the bioinformatics analysis. Yunqun Zhu reviewed and revised the manuscript. Yang Ji performed the formal analysis. Linxiang Lin conceived and designed the experiments and reviewed and edited the manuscript. All authors have read and agreed to the published version of the manuscript.

## Data Availability Statement

The data presented in this study are available in the article.

